# A nemertean excitatory peptide/CCHamide regulates ciliary swimming in the larvae of *Lineus longissimus*

**DOI:** 10.1101/634543

**Authors:** Daniel Thiel, Philipp Bauknecht, Gáspár Jékely, Andreas Hejnol

## Abstract

**Background:** The trochozoan excitatory peptide (EP) and its ortholog, the arthropod CCHamide, are neuropeptides that are only investigated in very few animal species. Previous studies on different trochozoan species focused on their physiological effect in adult specimens, demonstrating a myo-excitatory effect, often on tissues of the digestive system. The function of EP in the planktonic larvae of trochozoans has not yet been studied.

**Results:** We surveyed transcriptomes from species of various spiralian (Orthonectia, Nemertea, Brachiopoda, Entoprocta, Rotifera) and ecdysozoan taxa (Tardigrada, Onychophora, Priapulida, Loricifera, Nematomorpha) to investigate the evolution of EPs/CCHamides in protostomes. We found that the EPs of several pilidiophoran nemerteans show a characteristic difference in their C-terminus. Deorphanization of a pilidiophoran EP receptor showed, that the two isoforms of the nemertean *Lineus longissimus* EP activate a single receptor. We investigated the expression of EP in *L. longissimus* larvae and juveniles with customized antibodies and found that EP-positive nerves in larvae project from the apical organ to the ciliary band and that EP is expressed more broadly in juveniles in the neuropil and the prominent longitudinal nerve cords. While exposing juvenile *L. longissimus* specimens to synthetic excitatory peptides did not show any obvious effect, exposure of larvae to either of the two EPs increased the beat frequency of their locomotory cilia and shifted their vertical swimming distribution in a water column upwards.

**Conclusion:** Our results show that EP/CCHamide peptides are broadly conserved in protostomes. We show that the EP increases the ciliary beat frequency of *L. longissimus* larvae, which shifts their vertical distribution in a water column upwards. Endogenous EP may be released at the ciliary band from the projections of apical organ EP-positive neurons to regulate ciliary beating. A locomotory function of EP in *L. longissimus* larvae, compared to the association of EP/CCHamides with the digestive system in other animals suggests a dynamic integration of orthologous neuropeptides into different functions during evolution.

## Introduction

Neuropeptides are neurotransmitters that are involved in the regulation of most behavioral and physiological processes in animals. Many neuropeptide systems are ancestral to bilaterians and orthologous neuropeptides are deployed in the different bilaterian lineages independent of their nervous system organization (1–3). Only few of the orthologous neuropeptides are well conserved between the different bilaterian lineages and it is often the orthology of their receptors, that reveals their homology (1–4). For example, the protostome orthologs of vertebrate neuromedin -B/bombesin- and endothelin-related neuropeptides, the CCHamide and excitatory peptide (EP), are neuropeptides where the orthology of the deuterostome and protostome peptidergic systems could only be detected because of their receptor similarity (2, 3, 5).

The EP was initially discovered in the earthworms *Eisenia foetida* and *Pheretima vittata* (6) and has since been identified in many annelid and mollusk species where it has been described under various names depending on the taxon and due to either its myo-excitatory effect or its C-terminal structure (see table 1 for species, peptide names and references). The arthropod ortholog CCHamide was first discovered in the silk worm *Bombyx mori* (7) and is known from various arthropods, including insects (3, 7, 8), crustaceans (3, 8–10), myriapods (11) and chelicerates (8, 12, 13), with its nomenclature based on the presence of two conserved cysteine residues and an amidated C-terminal histidine residue. Due to the presence of the two cysteine residues, the usually amidated C-terminus and a similar precursor structure in CCHamides and EPs (Figure 1a), the two peptides were already recognized as possible orthologs (2, 3, 14) before the corresponding receptors were known. This orthology hypothesis was then confirmed with the deorphanization of their orthologues receptors (8, 15). Notably, the CCHamide system duplicated within the insect lineage into two distinctive CCHamides, which seem to specifically activate each their own receptor paralog (5, 8). Experiments showed that CCHamide is connected to feeding, sensory perception and the control of insulin like peptides in *Drosophila melanogaster* (16–18) and connected to feeding in other insects (5, 19). Its expression in different larval or adult insects is connected to the digestive system (7, 18–22). Experiments with EP showed a myo-excitatory effect that included digestive tissues of oligochaetes (6), leeches (23, 24), and a gastropod species (25), and association with digestive tissues was also shown by immunohistochemistry on a polychaete (26). (See also table 2 for functional and anatomical association of EP/CCHamides). The myo-excitatory effect and the expression of EP was observed on tissues of adult animals.

**Table 1:**
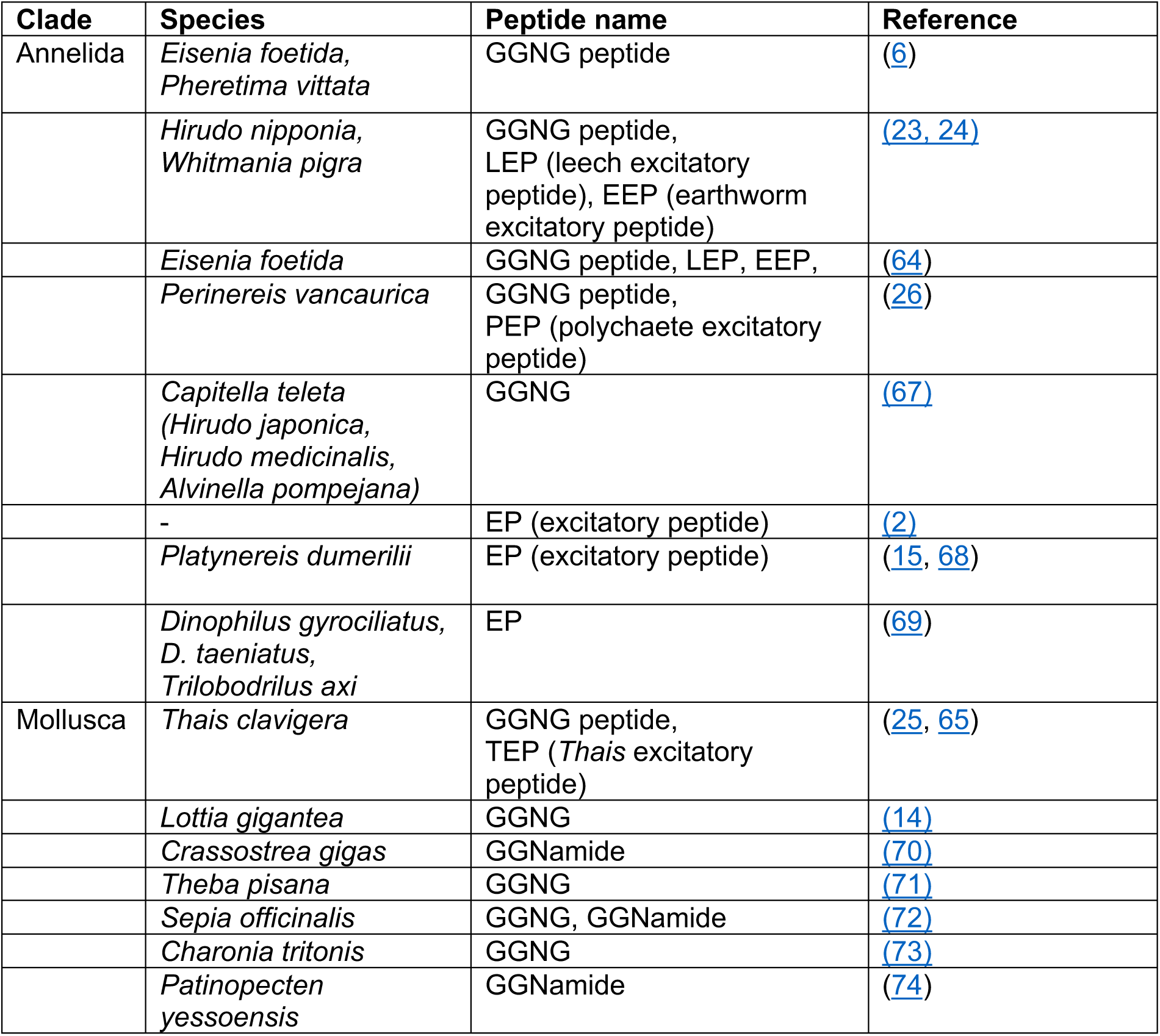
Discovery and nomenclature of trochozoan EPs.

**Table 2:** Association of EP/CCHamide peptidergic signaling based on expression, peptide detection and functional analysis of previous studies.

| Clade | Species | Results with regard to EP/CCHamide | Reference |
| --- | --- | --- | --- |
| Annelida | *Eisenia foetida*,<br>*Pheretima vittata* | Isolation of EP from gut tissue as well as whole bodies and excitation of gut tissue by EP application. | (6) |
|  | *Hirudo nipponia*, | Excitation of the crop gizzard by EP application. | (23) |
|  | *Eisenia foetida* | EP binding capacity is high in anterior part of digestive tract and the nephridia. | (64) |
|  | *Whitmania pigra* | EP immunoreactivity in supra-esophageal ganglion, circum-esophageal connective, sex segmental ganglion. | (24) |
|  | *Perinereis vancaurica* | EP immunoreactivity in CNS, epithelial cells of pharynx and epidermal cells. | (26) |
| Mollusca | *Thais clavigera* | Excitation of esophagus and penial complex by EP application, EP immunoreactivity in CNS and nerve endings of the penial complex. | (25) |
|  | *Thais clavigera* | EP1 expression in sub-esophageal, pleural, pedal and visceral ganglion and EP2 expression in pedal and visceral ganglion. | (65) |
| Hexapoda | *Bombyx mori* - larvae | CCHa expression in the central nervous system and the midgut. | (7) |
|  | *Phormia regina* - adults | CCHa2 injection stimulates feeding motivation (measured by the proboscis extension reflex at different sugar concentrations). | (5) |
|  | *Delia radicum* - larvae | CCHa1 was exclusively detected in the gut. | (21) |
|  | *Spodoptera exigua* - larvae | CCHa1 and CCHa2 are expressed in the larval gut and brain. Starvation increased CCHa1 expression in larvae. | (19) |
|  | *Drosophila melanogaster* - larvae & adults | High CCHa2 expression in gut and low expression in brain; high CCHa2 receptor expression in brain and low expression in gut. | (20) |
|  | *D. melanogaster* - adults | Upregulation of CCHa (1?) in the brain of starved animals.<br>RNAi knockdown of the CCHa1 receptor and CCHa1 receptor mutants showed an abolishment of a starvation-induced increase in olfactory responsiveness. | (16) |
|  | *D. melanogaster* - larvae & adults | Distinct CCHa1 and CCHa2 immunoreactivity in the digestive tract in both larvae and adults. | (22) |
|  | *D. melanogaster* - larvae | CCHa2 is highly expressed in fat body and slightly in gut, CCHa2 receptor is expressed in few endocrine cells in the brain including insulin like peptide (ILP) 2 producing cells. Starvation reduces CCHa2 expression. CCHa2 receptor mutants showed no change in ILP 2 and 3 expression but reduced ILP 5 expression. CCHa2 mutants show growth retardation and developmental delay. CCHa2 mutants show reduced ILP 5 expression and reduced body weight. | (18) |
|  | *D. melanogaster* - larvae & adults | Larvae: CCHa2 mutants show reduced feeding rate/activity and have a delayed development. Larvae and Pupae have reduced expression of insulin like peptide 2 and 3. CCHa2 is highly expressed in gut and slightly in brain. No effect detected for CCHa1.<br>Adults: CCHa2 mutants show reduced feeding and reduced locomotory activity. No effect detected for CCHa1. | (17) |
| Crustacea | *Marsupenaeus japonicus* – juvenile/adults | Highest expression of CCHa in ventral nerve cord, brain, eyestalks and gills, only low expression in intestines and stomach tissue. No effect of starvation on CCHa expression. | (66) |
|  | *Nephrops norvegicus* - adults | Tissue specific transcriptome detection of two CCHa's in brain, thoracic ganglia and eyestalks, but not in hepatopancreas or ovaries. | (10) |

**Figure 1:**
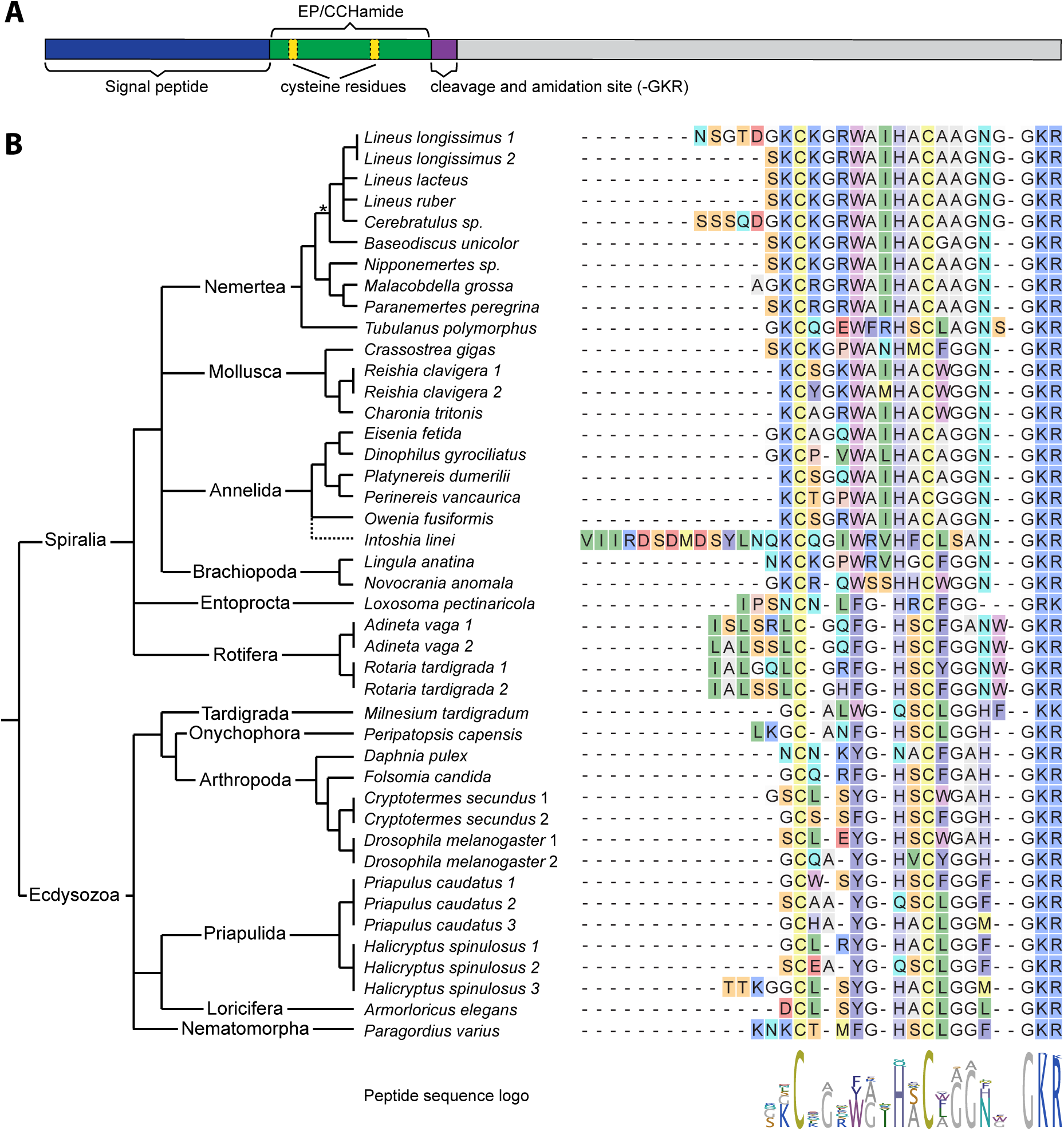
Protostome EP/CCHamide sequences. **A)** Schematic structure of EP/CCHamide precursors. **B)** Alignment of the predicted EP/CCHamide peptides of different protostomes with the phylogenetic relationship of the different taxa. C-terminal GKR, GKK or GRK residues indicate the precursor cleavage site and a C-terminal amidation of the residue N-terminal to the glycine, a missing glycine residue (e.g. *M. tardigradum*) indicates only cleavage without amidation. Peptide sequence logo was created from the alignment. Phylogeny is depicted according to Dunn et al. 2014 (28), annelid phylogeny according to Struck et al. 2015 (55) with Orthonectida as an annelid taxon (56), arthropod phylogeny according to Yeates et al. 2016 (57) and nemertean phylogeny according to Andrade et al 2014 (58) and Kvist et al. 2015 (59). Dashed line indicates unclear relationship, asterisks indicate the heteronemertean branch. Full precursor sequences are available in the Supplementary Material.

Annelids and molluscs, as well as other closely related taxa such as brachiopods and nemerteans, also possess planktonic larvae that usually metamorphose into morphologically different, benthic adults. This stage called ‘trochophora larva’ is the name giving characteristic of the spiralian taxon that is referred to as Trochozoa (27–31). While antibodies against neuropeptides (usually FMRFamide) have been used widely to describe neuroanatomies of trochozoan larvae (32–40), there are comparably few studies that investigated the behavioral effect of neuropeptides in such larvae (41–45) and neither behavioral nor immunohistochemical studies investigated the EP/CCHamide in trochozoan larvae.

Here we report about the evolution of EP/CCHamide orthologs in ecdysozoans and spiralians with a focus on nemerteans. We test the activation of a single EP receptor by two EP isoforms in the nemertean *L. longissimus*, the EP expression in the pilidium larvae and juveniles of *L. longissimus*, and the influence of the EP on the behavior of *L. longissimus* larvae.

## Material and Methods

### Bioinformatics

Publicly available transcriptomic or genomic sequencing data were scanned for EP precursors or EP receptors: Adineta vaga (Ensembl: AMS_PRJEB1171), Armorloricus elegans (NCBI: SRR2131253), Baseodiscus unicolor (NCBI: SRR1505175), Cerebratulus spec. (NCBI: SRR1797867), Dinophilus gyrocilatus (NCBI: SRR4039542), Halicryptus spinulosus (NCBI: SRR2682062), Intoshia linei (NCBI: PRJNA316116), Lineus lacteus ((46) dryad data package), Lineus longissimus (NCBI: SRS1118155), Lineus ruber ((46) dryad data package), Loxosoma pectinaricola ((46) dryad data package), Malacobdella grossa ((46) dryad data package), Milnesium tardigradum (NCBI: SRR1598869), Nipponemertes spec. (NCBI: SRR1508368), Novocrania anomala (NCBI: SRS1118148), Owenia fusiformis (NCBI: SRS590031), Paragordius varius (NCBI: PRJEB19315), Paranemertes peregrine ((46) dryad data package), Peripatopsis capensis (NCBI: SRR1145776), Priapulus caudatus (NCBI: SRR1212719), Rotaria tardigrade (NCBI: SRR2430032), Tubulanus polymorphus ((46) dryad data package).

Publicly available references sequences from NCBI and from publications that are listed in Table 1 and Table 2 were used as BLAST reference sequences. Neuropeptide precursor sequences were tested for the presence of a signal peptide using SignalP 4.1 (47) and Signal-3L 2.0 (48). Receptor sequences were aligned with ClustalX v2.1 (49), non-conserved regions where removed with TrimAl (using the gappyout option) (50), the phylogenetic tree was calculated with FastTree v2.1 (51) (using the LG amino acid substitution model) and visualized with FigTree v1.4.3 (http://tree.bio.ed.ac.uk/software/figtree).

### Receptor activation assay

For the receptor activation assay the protocol of (15) was followed. The full ORF of the *L. longissimus* EP-receptor sequence was amplified by PCR from larval cDNA and was cloned into the mammalian expression vector pcDNA3.1(+). The vector was transfected into CHO-K1 cells together with a promiscuous G_α_-16 protein encoding plasmid and a calcium sensitive luminescent apoaequorin-GFP fusion protein encoding plasmid (G5A). Synthetic peptides (customized from GenScript) were diluted to different concentrations, the dose-dependent luminescence response was measured in a plate reader (BioTek Synergy Mx and Synergy H4, BioTek, Winooski, USA) and analyzed with Prism 6 (GraphPad, La Jolla, USA).

### Animal rearing

Adult *Lineus longissimus* (Gunnerus, 1770) and *Novocrania anomala* (O. F. Müller, 1776) were collected at different collection sites close to the Marine Biological Station Espegrend (University of Bergen) in Norway. *L. longissimus* were kept together at 12°C in natural seawater until spawning occurred naturally during spring. Larvae were kept in 5-liter beakers with slow rotators and were fed *Rhinomonas* sp. algae.

### Immunohistochemistry

Antibodies against the C-terminal CAAGNGamide sequence of the *Lineus* excitatory peptide were raised in rabbits (genscript) and the antibodies were cleaned from the blood serum with a Sulfolink™ Immobilization Kit (ThermoFisher Scientific) according to (32). *L. longissimus* larvae were relaxed in an 8% MgCl_2_*6H_2_O solution in distilled water for 10 minutes, fixed in a 4% Formaldehyde solution in seawater for 1 hour, washed 5*5minutes in PBS + 0.1% Tween (PTw), transferred into methanol and stored at -20°C. After rehydration in PTw, specimens were incubated in PBS + 0.5% Triton X for 4h at room temperature, transferred into a 0.1M Tris pH 8.5 + 0.1% Tween solution (THT), blocked for 1 h in a 5% normal goat serum (NGS) solution in THT and incubated with the primary antibody in THT + 5% NGS for 3 days (approximately 1 ng/µl anti-EP antibody from rabbit and 0.5 ng/µl anti tyrosinated tubulin antibody from mouse). The primary antibodies were washed 2*5 min in THT containing 0.9 M NaCl, washed 5*15 min in PBS + 0.2% Triton X + 0.1% BSA (PBT), then washed 5*30 min in PTw, transferred into THT, blocked for 1h in THT + 5% NGS and incubated with the secondary antibodies in THT + 5% NGS overnight (Alexa 488 anti rabbit and Alexa 647 anti mouse). Specimens were then washed 5*15 min in PBT, 2*30 min in PTw, incubated 1h in PTw containing 1 µg/ml DAPI, washed 2*30 min in PTw and transferred into 70% glycerol before mounting in 70% glycerol. As a control to test the antibody specificity, antibodies were preabsorbed in a 2 mM solution of synthetic full-length EP in THT + 5% NGS during the blocking step of the primary antibody.

### Vertical swimming assay

Late larvae of *L. longissimus* (2-3 week old) were recorded in transparent columns (6.5 cm height / 3.7 cm length / 2.2 cm width) to compare the vertical distribution of EP-exposed and untreated larvae. For each treatment and replicate more than 200 larvae per column were recorded with a “DMK31AU03” camera (The Imaging Source ®) with LED illumination in a darkened box to avoid uncontrolled effects due to the possibility of photo-behavior of the larvae. All treatments and recordings were repeated twice, each time with treatment and control being recorded next to each other. The videos were processed and analyzed with “Fiji” (52) and the repeats were averaged afterwards. (Fiji measurements: 1. Background elimination using Z-project (Average Intensity of all frames) and Image calculator (substract Z-projection from all frames of the video), 2. Conversion of moving larvae to black dots using individual threshold parameter, 3. Combining distribution of larvae over time intervals of 5-10 seconds using Z-project (Standard Deviation of the corresponding frame sequences), 4. Measurement of larval distribution by dividing the height of the column into 50 squares and using the ‘Measure’ function on each height division, 5. Comparing the measurements in an table to get a horizontal profile of distribution of larvae.) The distributions were compared and tested with a two sample Kolmogorov-Smirnov test.

### Measurement of the ciliary beat frequency of *L. longissimus* larvae

First attempts to immobilize *L. longissimus* larvae between a microscopic glass slide and cover slip failed, as the larvae were either slowly moving away or they got stuck and the cilia that touched the glass stopped beating rhythmically. Therefore, pulled glass holding-capillaries with an opening of about 50-70 µm were used to immobilize the pilidium larvae, similar to (53). Larvae were caught with the capillaries at their apical tip between a glass slide and a cover slip that was completely filled with seawater. Reagents were added on one side of the cover slip with a pipette and simultaneously soaked off from the other side with a tissue paper, enabling paired measurements of the ciliary beating from individual larvae, under control conditions, after soaking in peptides and after peptide washout in. The ciliary beating was recorded with a DMK23UV024 camera (The Imaging Source) with 50 frames per second.

Preliminary measurements were taken to determine the larval excitation that was triggered by being immobilized and sucked to a holding capillary or by the water flow from changing the solutions. The tests showed that the ciliary beating slowly decreased after larvae were caught with the holding capillary, reaching after about ten minutes a point where the beat frequency did not further decrease. The next test showed that an initial increase in the ciliary beat frequency caused by an exchange of the solution decreased back to the normal frequency in less than two minutes. Test with several exchanges in frequent intervals showed that the ciliary beating was always back to the initial frequency after less than two minutes. Based on these preliminary tests, measurements were taken as follow: Larvae were caught with a glass capillary, the slide got transferred to the microscope and after about ten minutes the seawater was exchanged with fresh seawater twice in a row with a waiting time of two minutes after each exchange, before the control condition was recorded. The seawater solution was then exchanged a third time with seawater containing one of the peptides, and after two minutes the ciliary beating under the influence of EP was recorded. Afterwards the larvae were washed for 15 minutes with several exchanges of fresh seawater before the ciliary beating for the washout was recorded. Generally, the beating of apical and lateral ciliary bands was tested for 12 larvae under the influence of EP1 and 12 larvae under the influence of EP2, measuring control, peptide exposed and washout conditions for every larva. The videos were processed with Fiji (“plot Z-axis Profile” function to measure the frequency), and differences in the ciliary beat frequency were tested for significance with a paired t-test.

## Results

### Transcriptome analyses show a wide distribution of EP/CCHamide orthologs in protostomes and an extended C-terminus in several heteronemerteans

We identified spiralian orthologs of EP/CCHamide in transcriptomes of nemertean, brachiopod, entoproct and rotifer species, and ecdysozoan orthologs in tardigrade, onychophoran, priapulid, loriciferan and nematomorph species, showing the wide distribution of EP/CCHamide within protostomes (Fig. 1). Nearly all peptides possess two conserved cysteine residues, an amidated C-terminus and a propeptide structure with a single copy of the bioactive peptide directly located after the signal peptide (Fig. 1, Supplementary list of precursors). The main exception seems to be the EP ortholog in the tardigrade *Milnesium tardigradum*, which lacks the C-terminal amidation and also has a precursor where the signal peptide and the EP/CCHamide peptide are separated by several amino acids – although a similar precursor structure has also been described for the leech *Hirudo nipponia* (54) and the CCHamide 2 of *D. melanogaster* (5). The name-giving C-terminal histidine residue of the arthropod CCHamide is only present in Arthropoda and Onychophora. This residue seems more variable within ecdysozoans, with methionine, phenylalanine or leucine residues in priapulid, nematomorph and loriciferan species. All investigated spiralians share an asparagine residue in this position, although while in most taxa this seems to be the most C-terminal residue, both investigated rotifer species and several nemerteans possess an extended C-terminus with an additional amino acid. The closely related heteronemertean *Cerebratulus* and *Lineus* species all possess an additional glycine residue. This glycine residue seems to have evolved within the heteronemertean lineage, as neither the more distantly related heteronemertean *Baseodiscus unicolor*, nor the non-heteronemerteans showed this additional residue.

### Two *L. longissimus* excitatory peptides activate a single *L. longissimus* EP receptor

In the nemertean *L. longissimus* we identified two excitatory peptide transcripts. The peptides differed only in a few amino acids at the N-terminus of the predicted EP/CCHamide peptide. PCRs using larval cDNA and two specific primer pairs that were designed in the region where the two transcripts vary (Supplementary Material) confirmed the presence of two separate transcripts. We only identified a single ortholog of EP/CCHamide receptors in the transcriptome of *L. longissimus*. A phylogenetic analysis shows the orthologues relationship of protostome EP and CCHamide receptors and the chordate neuromedin B/bombesin and endothelin related receptors (Fig. 2 A). The analysis also shows the duplication of the CCHamide receptors in the early insect lineage, which is not present in Collembola, but in as distantly related pterygote insects as dipterans and termites. This receptor duplication within pterygote insects and its absence in Collembola is in accordance with the presence of two CCHamide paralogs in pterygote insects (Fig. 1 B).

**Figure 2:**
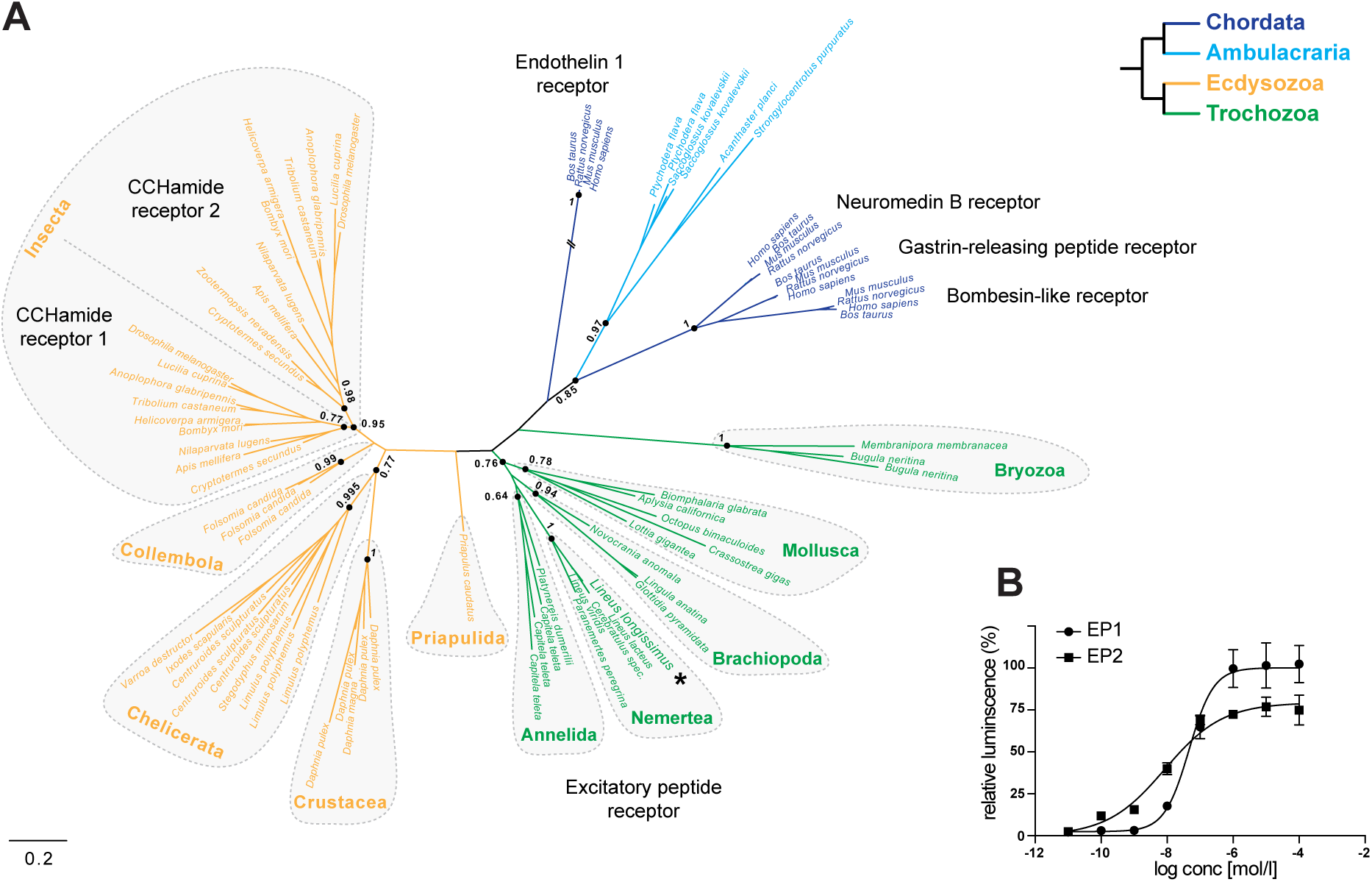
Analysis of EP/CCHamide receptors. **A)** Phylogenetic analysis of protostome EP/CCHamide receptors and related deuterostome receptors. Color coding according to the phylogenetic group as depicted in the simplified tree on the upper right corner. SH-like support values are shown for the indicated nodes. Scale bar on the lower right corner shows amino acid substitution rate per site. The endothelin 1 receptor branch was shortened to half its size (indicated by the two crossing lines). Asterisk highlights the *L. longissimus* receptor that was biochemically characterized. **B)** Activation of the *L. longissimus* EP receptor by the two *L. longissimus* excitatory peptides EP1 and EP2.

We cloned the *L. longissimus* EP receptor into an expression vector, expressed it in mammalian cells and tested its activation by the *L. longissimus* EP1 and EP2 peptides. Both peptides activated the receptor in a similar, nano-molar concentration with EC_50_ values (half maximal effective concentration) of 78 nmol/l for EP1 and 59 nmol/l for EP2 (Fig. 2 B).

### EP is expressed in neurons that project from the apical organ to the ciliary bands of *L. longissimus* larvae

We visualized EP expressing neurons in *L. longissimus* pilidium larvae using polyclonal antibodies against the C-terminal residues CAAGNGamide. The larval staining revealed EP positive neurons in the apical part of the larvae as well as in developing juveniles inside the larvae (Fig. 1 A). The larvae showed the apical EP-positive nerves already in 5-day old young larvae before development of the juveniles. The EP positive nerves of the larvae project from the lateral organ towards the ciliary bands at the anterior end of the future juvenile (Fig. 3 B, C). The antibodies stained axons as well as somata of a variable number of cells (Fig. 3 C, D), possibly depending on the age of the larva. The projecting EP-positive neurons directly innervate the prominent neurons underneath the ciliary band at the point where the ciliary bands of the apical lobes and lateral lappets meet (Fig. 3 E). With strong excitation and sensitive detection setting, the nerves of the ciliary bands themselves also showed weak EP immunoreactivity (Fig. 3 F, G), however, the intensity of the labelling was weaker than the one of the nerves that project from the apical organ and we could not detect EP positive cell bodies that might belong to the ciliary nerve. The EP positive neurons of the larvae and the developing juveniles are not connected. Hatched juveniles show prominent EP staining in the neuropil and in the two longitudinal ventral nerves along the whole body (Fig. 3 H, I).

**Figure 3:**
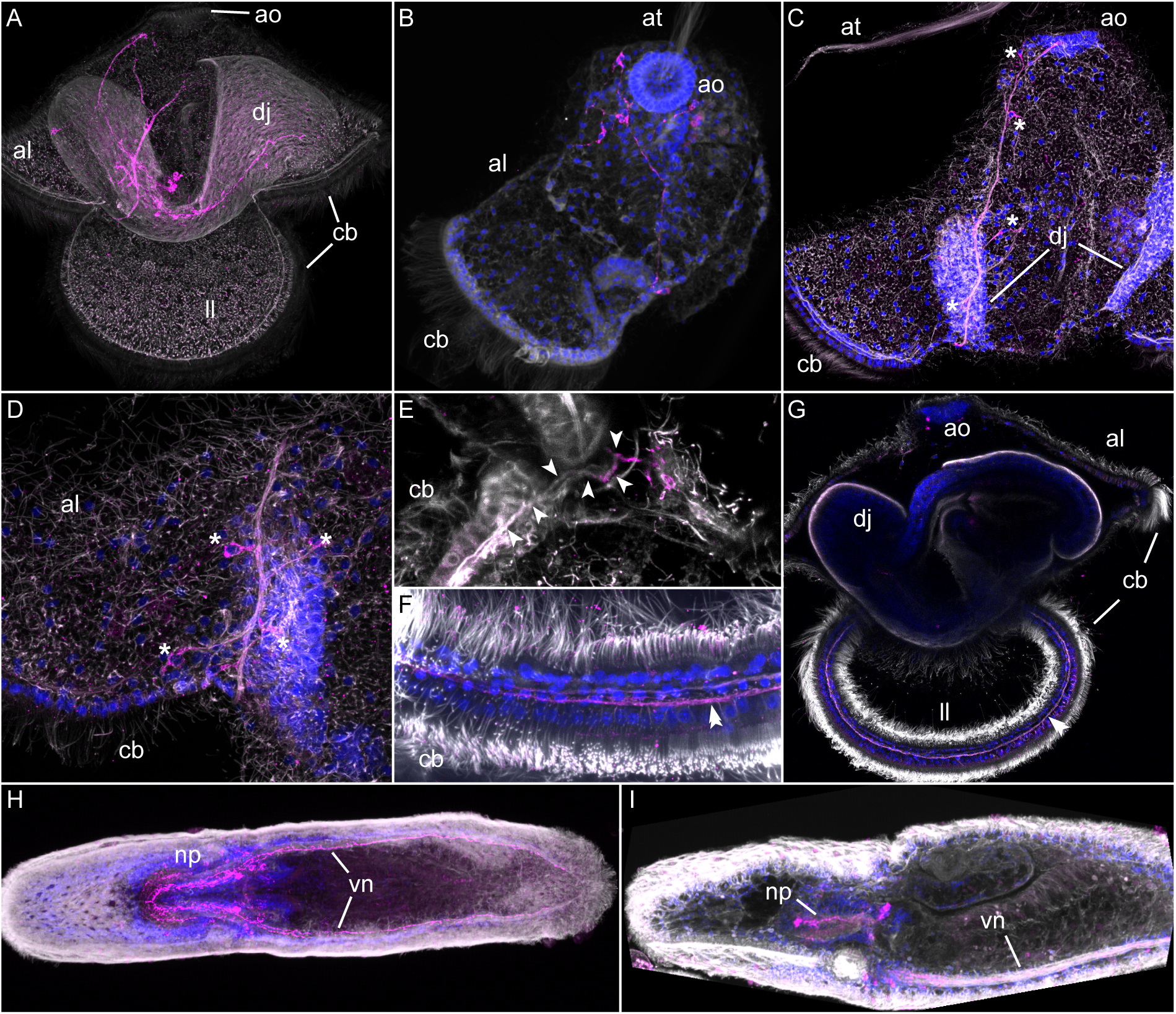
Immunohistochemical analysis of EP in *L. longissimus* larvae and juveniles. **A)** Side-view of a larva with advanced developing juvenile, showing nerve underneath the ciliary bands and the EP positive nerves in the apical lobe and in the juvenile. **B)** Top-view of the apical part of a larva, with apical organ and ciliary band of the apical lobe, showing network of EP positive nerves. **C)** Side-view of the apical part of a larva, showing EP positive nerve running from the apical organ towards the ciliary band. **D)** Side-view of the EP positive nerve before the innervation of the ciliary band, including EP positive cell bodies of the nerve cells. **E)** Innervation of the nerve underneath the ciliary band by a more proximal EP-positive part. Arrows follow the nerve with strong proximal EP signal in the beginning. **F)** Close-up of EP positive signal in the nerve underneath the ciliary of a lateral lappet after strong signal amplification. **G)** Side-view of a larva with developing juvenile, showing EP positive signal in the nerve underneath the ciliary after strong signal amplification. Double arrow indicates the nerve. **H)** Top-view of a juvenile with broad EP-positive signal in the neuropil and the ventral nerve cords. **I)** Side-view of a juvenile with broad EP-positive signal in the neuropil and the ventral nerve cord. *Orientation*: all larval pictures are oriented with the anterior side of the developing juvenile to the left. *Abbreviations*: al = apical lobe, ao = apical organ, at = apical tuft, cb = ciliary band, dj = developing juvenile, ll = lateral lappet, np = neuropil, vn = ventral nerve cord. *Color coding*: gray/white = anti-tyrosinated tubulin staining (cilia and nerves), magenta = anti-EP staining (Ep positive nerve cells), blue = DAPI staining (nuclei).

### EP influences the swimming behavior of *L. longissimus*

We tested the influence of the predicted EP1 and EP2 peptides on *L. longissimus* specimens by soaking them with synthetic peptides. When we exposed about 3-week-old planktonic pilidium larvae to 5 µmol/l EP1 or EP2 and recorded their swimming behavior in a vertical column, their average swimming level was shifted upwards, and larvae concentrated in the upper part of the water column (p = 3.3e-3 for EP1 and 6.6e-4 for EP2) (Fig. 4 A). The vertical distribution of larvae exposed to EP1 and EP2 seemed similar (p = 0.94) and after washout of the peptides, the formerly treated larvae returned to a similar distribution as washed out control larvae (p = 0.95 for EP1-washout/Ctrl-washout and 0.82 for EP2-washout/Ctrl-washout). Juveniles did not show any obvious reaction at any tested concentration between 50 nmol/l and 50 µmol/l.

**Figure 4:**
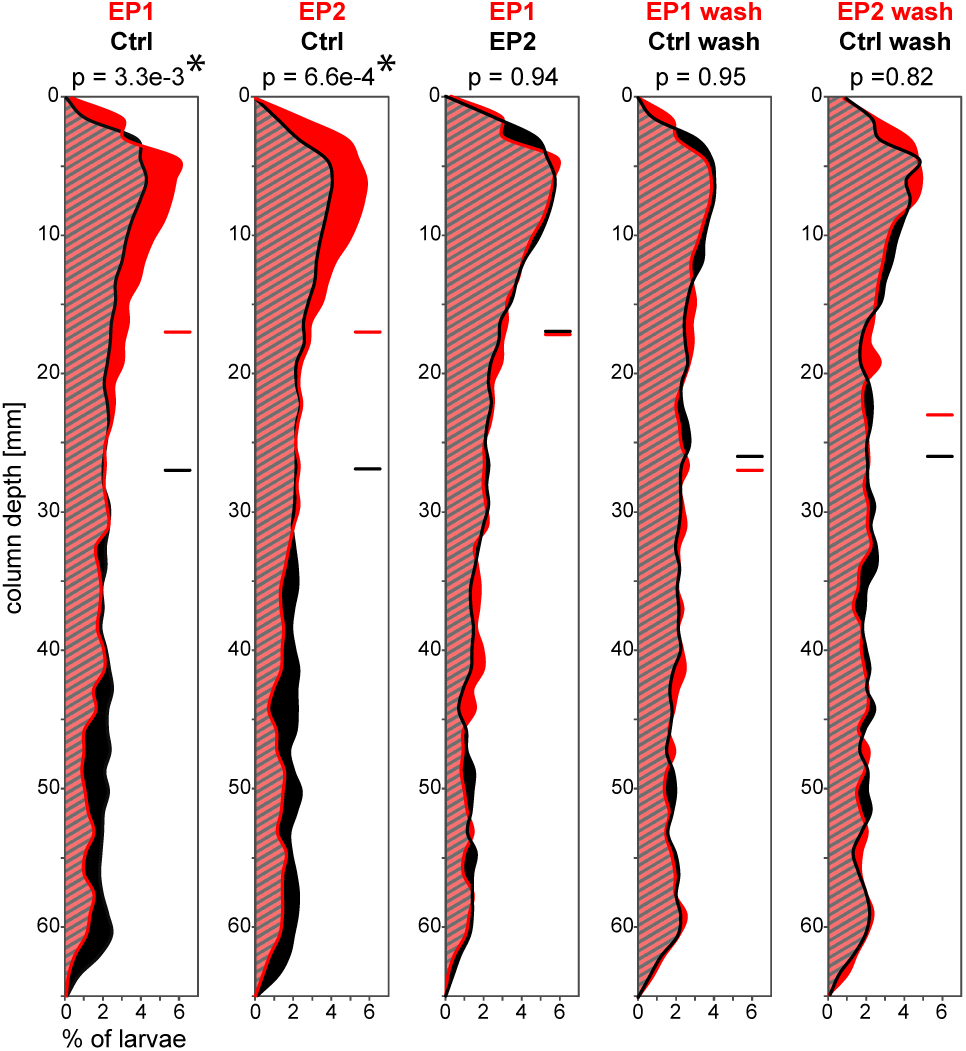
Influence of EP1 and EP2 on the vertical distribution of *L. longissimus* larvae in a water column. Comparisons from left to right: control and EP1 exposed specimens, control and EP2 exposed specimens, EP1 and EP2 exposed specimens, control and EP1 exposed specimens after several water changes, control and EP2 exposed specimens after several water changes. Y-axis shows water depth of column, X-axis shows the percentage of larvae. Red and black lines show distribution of larvae under the condition stated above the columns. Horizontal lines indicate average swimming height. P values compare the distribution of the two samples (two-sample Kolmogorov-Smirnov test). Ctrl = control, wash = washout.

### EP leads to an increase in the beat frequency of apical and lateral ciliary bands of *L. longissimus* larvae

To test whether the upwards shift of the swimming distribution of *L. longissimus* larvae after exposure to EP may be due to a change in the beat frequency of their locomotory cilia (Fig. 5 A), we recorded and quantified the beat frequency of locomotor cilia before, during and after exposure to excitatory peptide (Fig. 5 B,C). When larvae were exposed to concentrations of 25 µmol/l or higher of EP1 or EP2, the cilia stopped beating rhythmically and started standing up straight instead, while vibrating with a high frequency (Fig. 5 D, E). At lower concentrations, we detected a significant increase of the ciliary beat frequency (cbf) of the outer ciliary bands of the lateral lappets and apical lobes (Fig. 5 F). EP1 seemed to be less efficient than EP2, as 10 µmol/l EP1 were necessary to observe significant changes in the cbf, compared to only 5 µmol/l EP2. When the peptides were washed out, the cbf decreased again. We did not determine a possible effect of EP on ciliary arrests, as such arrests only occurred in combination with muscular contractions of the lobes and are likely not associated to locomotion, but rather a mechano-sensory response that is related to feeding (53). These contractions also seemed to be rather irregular and increased with the time the larvae were stuck to the holding pipette. The ciliary bands of the lateral lappets showed under control conditions a significantly higher cbf than the ciliary bands of the apical lobes with an average of 10.34 beats per second (bps) and 9.66 bps, respectively (p = 1.17e-7, n = 24). Exposure to EP1 or EP2 cancelled this difference, so the cbf of the lateral and apical ciliary bands became more similar with 10.98 bps and 10.79 bps, respectively (p = 0.083, n = 24; see Fig. 5 F for differences between EP1 and EP2 treated animals).

**Figure 5:**
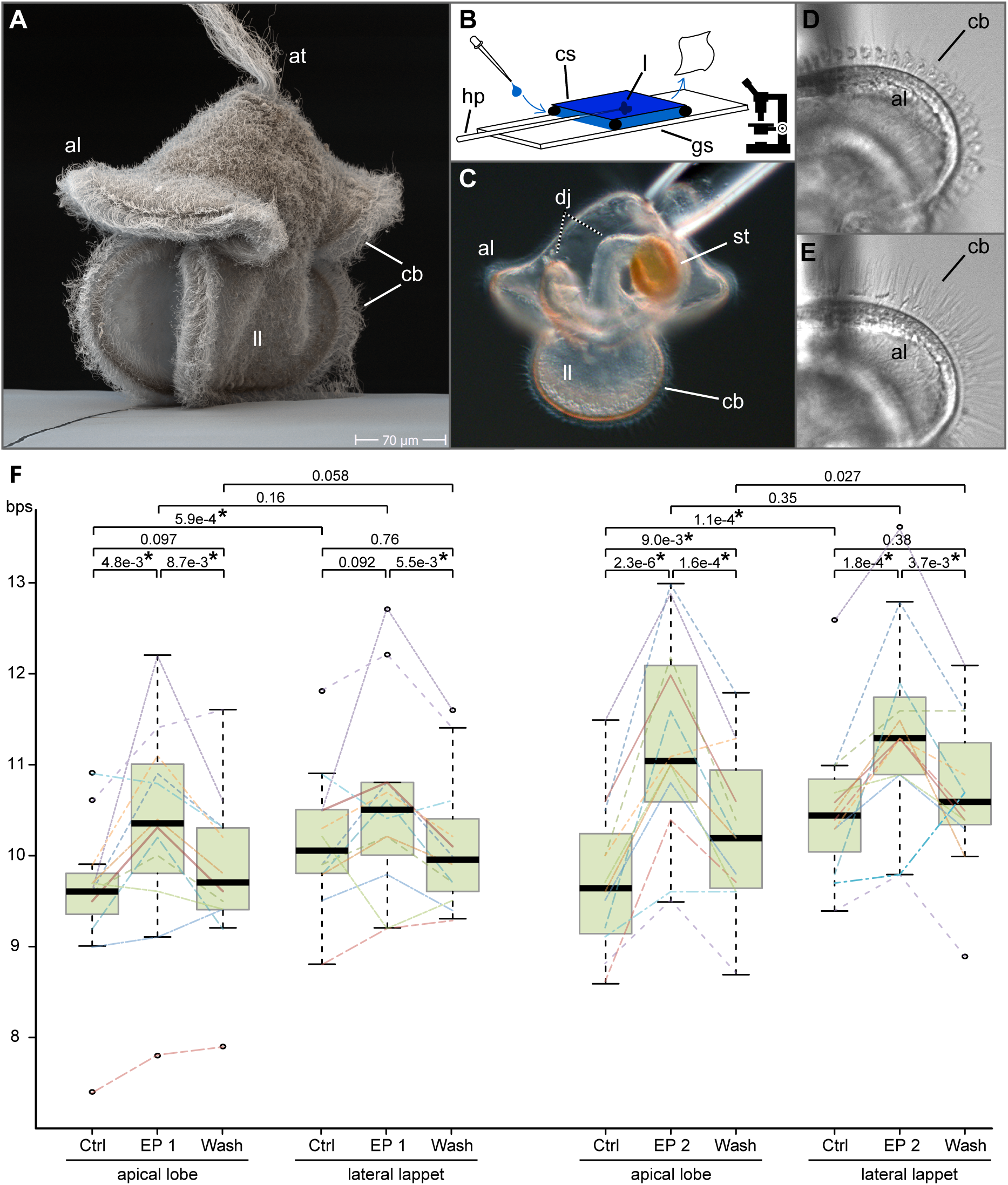
*L. longissimus* pilidium larvae and the influence of excitatory peptide on their ciliary beating. **A)** SEM picture of a *L. longissimus* larva. **B)** Experimental setup to record the ciliary beating and the influence of excitatory peptide on a microscope slide. **C)** Light microscopic picture of *L. longissimus* larvae with a developing juvenile on a holding pipette. **D)** Beating ciliary band of the apical lobe. **E**) The same ciliary band as in D, after overexposure to 20 µmol/l EP2. **F)** Boxplot of the ciliary beat frequency of the apical lobes and lateral lappets under normal conditions, after exposure to EP, and after peptide washout. Bold horizontal line indicates median, box indicates upper and lower quartile, whiskers indicate variability outside the upper and lower quartile, circles indicate outliers. Lines within the graph indicate measurement points of individual larvae. P values were calculated with a paired t-test. Asterisks indicate significance with p < 0.02. al = apical lobe, at = apical tuft, bps beats per second, cb = ciliary bands, cs = cover slip, Ctrl = control, EP = excitatory peptide, gs = microscopic glas slide, hp = holding pipette, dj = developing juvenile, l = larva, ll = lateral lappet, st = stomach.

## Discussion

### Reconstruction of the ancestral EP/CCHamide

When reconstructing the sequence of the ancestral protostome EP/CCHamide, a few characteristics can be assumed: The ancestral EP/CCHamide peptide likely possessed two cysteine residues, an amidated C-terminus and a propeptide structure where the signal peptide is directly followed by a single copy of the predicted ligand. Nearly all EP/CCHamide orthologs showed either two consecutive glycine residues or a glycine and an alanine residue between the second cysteine and the amidated C-terminus of the predicted ligand. Other features that seem to be shared are the presence of an aromatic amino acid and a histidine residue between the two conserved cysteine residues. A different peptide that might be related to the EP/CCHamide/neuromedin B peptides is elevenin (L11). This peptide is so far only known from protostomes and has a similar propeptide structure as EP/CCHamide and possesses two cysteine residues (2, 60). The relationship of the L11 and EP/CCHamide/neuromedin B receptors, however, has only been weakly supported in phylogenetic analysis (60) or has not been recognized at all (1, 15). If these peptidergic systems are indeed ancient paralogs, then at least the cysteines and the propeptide structure seem to be characteristics of their common ancestor. A similar propeptide structure with a single N-terminal peptide is also characteristic for the related deuterostome neuromedin B and gastrin-releasing peptides, both of which are also amidated at their C-terminus (3, 61). Cysteine residues, however, are only present in endothelins, but those peptides possess four cysteines instead of two (62, 63). The only true indication for an orthology of these deuterostome peptides with the protostome EP/CCHamides are - as previously shown - their receptors (1–3), and it is difficult to propose more ancestral characteristics than the described propeptide structure with a single N-terminal peptide and likely an amidated C-terminus - both of which can be found in at least one of the paralogous systems in the deuterostome lineage an the protostome lineage.

### Peptidergic nerves that influence ciliary swimming project from sensory organs to the ciliary band

EP-positive nerves in *L. longissimus* larvae project from the apical organ and innervate the nerves underneath the ciliary bands of the apical and the lateral lappets at the point where the lateral lappets and apical lobe merge. The EP positive nerve seems to directly connect sensory input and locomotory output organ. The immunoreactivity in the nerve of the ciliary band of *L. longissimus* was only detectable with strong excitation and sensitive detector settings, and it is not clear whether this signal can be attributed to possible background signal or very low expression levels. In larvae of the annelid *P. dumerilii*, most peptides which influence the ciliary based swimming can be found in the ciliary ring nerve (41). Many of these peptides in *P. dumerilii*, show immunoreactivity in apical organ sensory cells as well. In the veliger larvae of the mollusk *Crepidula fornicata*, FMRFamide showed to influence the ciliary beating and the larvae show strong FMRFamide-like immunoreactivity in the velar lobes projecting from the stalk towards the ciliary band and strong immunoreactivity in the nerves underneath the ciliary band (43). In the brachiopod larvae of *Terebratalia transversa*, the FMRFamide orthologue FLRFamide also influences the ciliary based swimming and FLRFamide-positive nerves project from the central neuropil underneath the apical organ towards the ciliary band, but no signal is detected underneath the ciliary band itself (44). In summary, it seems like nerve cells with peptides that can influence the ciliary based swimming often connect sensory organs and ciliary bands. Although immunoreactivity of those peptides can persist in the ciliary nerve band itself, this is not a general rule that applies to all such peptides.

### Functional association of peptides can vary during evolution

There are only few studies that test the behavioral effect of neuropeptides on trochozoan larvae (41–45) and those focus on peptides other than the EP/CCHamide. One of these studies shows that a variety of different neuropeptides can influence the ciliary beating in a single species (41). The behavioral effect of a single type of peptide on different animals has only been tested for FMRFamide and its orthologue FLRFamide and the different studies showed some species-specific effects in different trochozoan larvae (41, 43–45). The influence of EP on locomotion of *L. longissimus* larvae and the broad expression in the brain and prominent ventral nerve cord of the juveniles stands in contrast to the myo-excitatory effect of EPs in different adult/juvenile annelid (6, 23, 24, 26) and mollusc (25) species and the repeated association of EPs and CCHamides with feeding or the digestive system in annelids (6, 23, 24, 26, 64), molluscs (25, 65) and insects (5, 7, 16–22) (see also table 2). This suggests that the degree of functional conservation can vary depending on the type of peptide, the animal taxon or the developmental stages and that also functionally more conserved peptides like the EP/CCHamides can show plasticity in their behavioral association. This is for example also reflected in the lack of food and digestive tract association of CCHamide in the kuruma shrimp *Marsupenaeus japonicus* (66). A bigger comparison of the effect of EPs on larvae from other trochozoan taxa could give more insight into the conservation of the here described effect of EP/CCHamide orthologues within trochozoan larvae.

## Acknowledgements

We want to thank Carmen Andrikou and Anlaug Furu Boddington for helping with animal collection and rearing. We further thank Jürgen Berger from the Max Planck Institute for Developmental Biology (Tübingen, Germany) for taking the SEM picture of the larva. This research was supported by the FP7-PEOPLE-2012-ITN grant no. 317172 “NEPTUNE” and received further support by the DFG - Deutsche Forschungsgemeinschaft to GJ (Reference no. JE 777/3-1).

## Supplementary Material

All Data and Supplementary Material is available from the authors upon reasonable request.

## Abbreviations used in the text

bps: beats per second
cbf: ciliary beat frequency
EP: excitatory peptide
EC_50_: half maximal effective concentration
*D. melanogaster*: *Drosophila melanogaster*
*L. longissimus*: *Lineus longissimus*
L11: elevenin

## Supplementary: EP/CCHamide precursor sequences

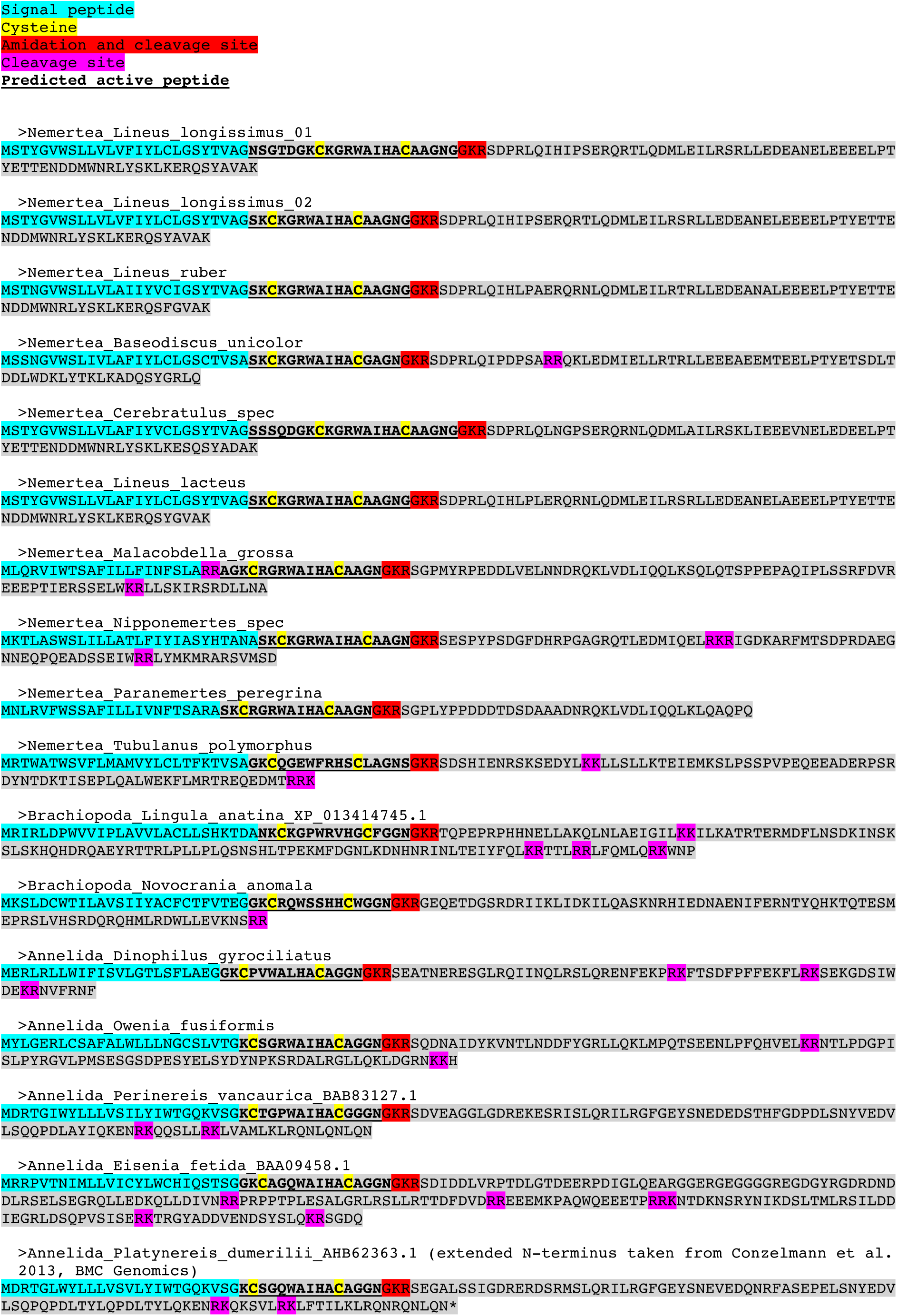
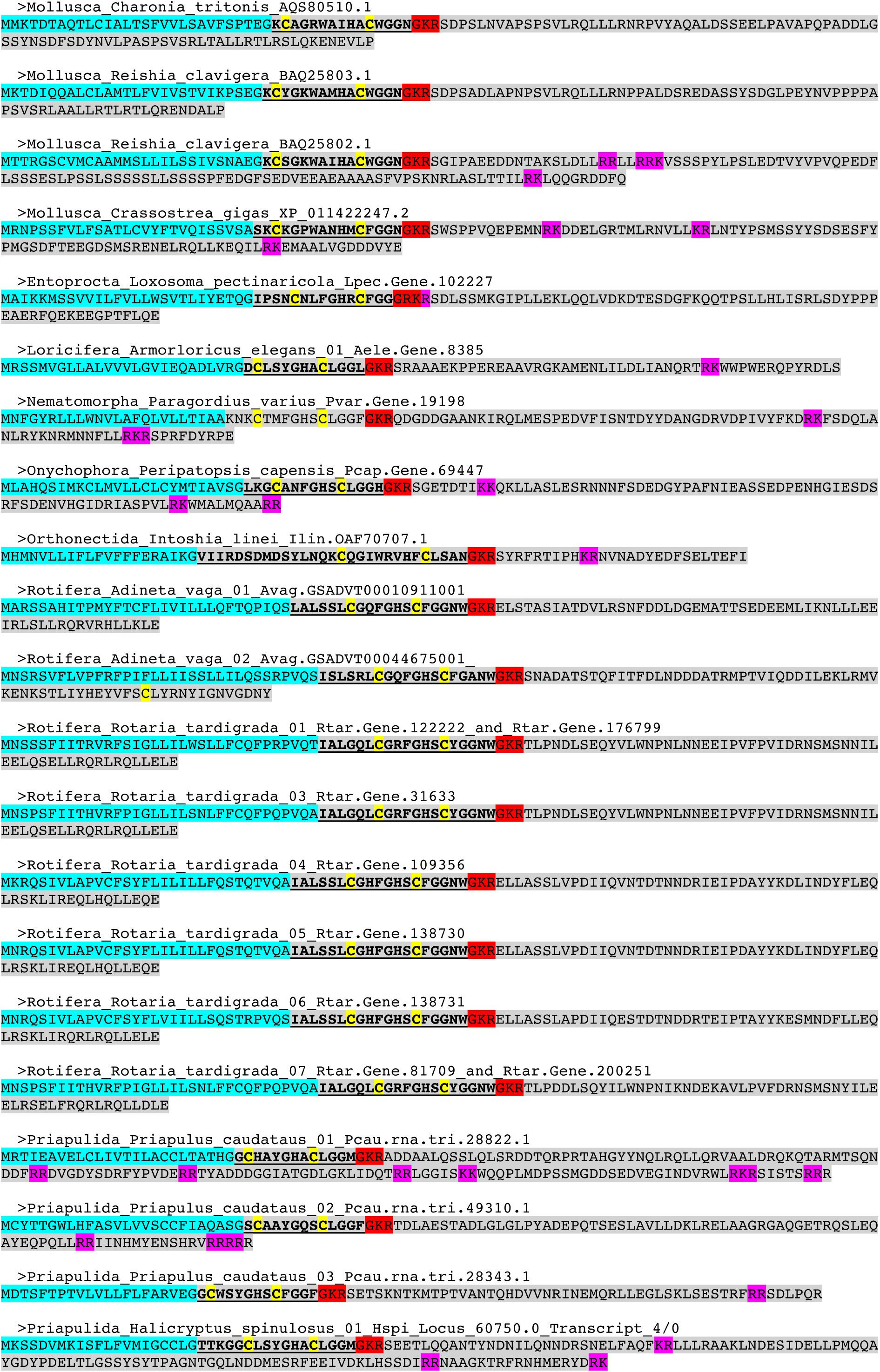
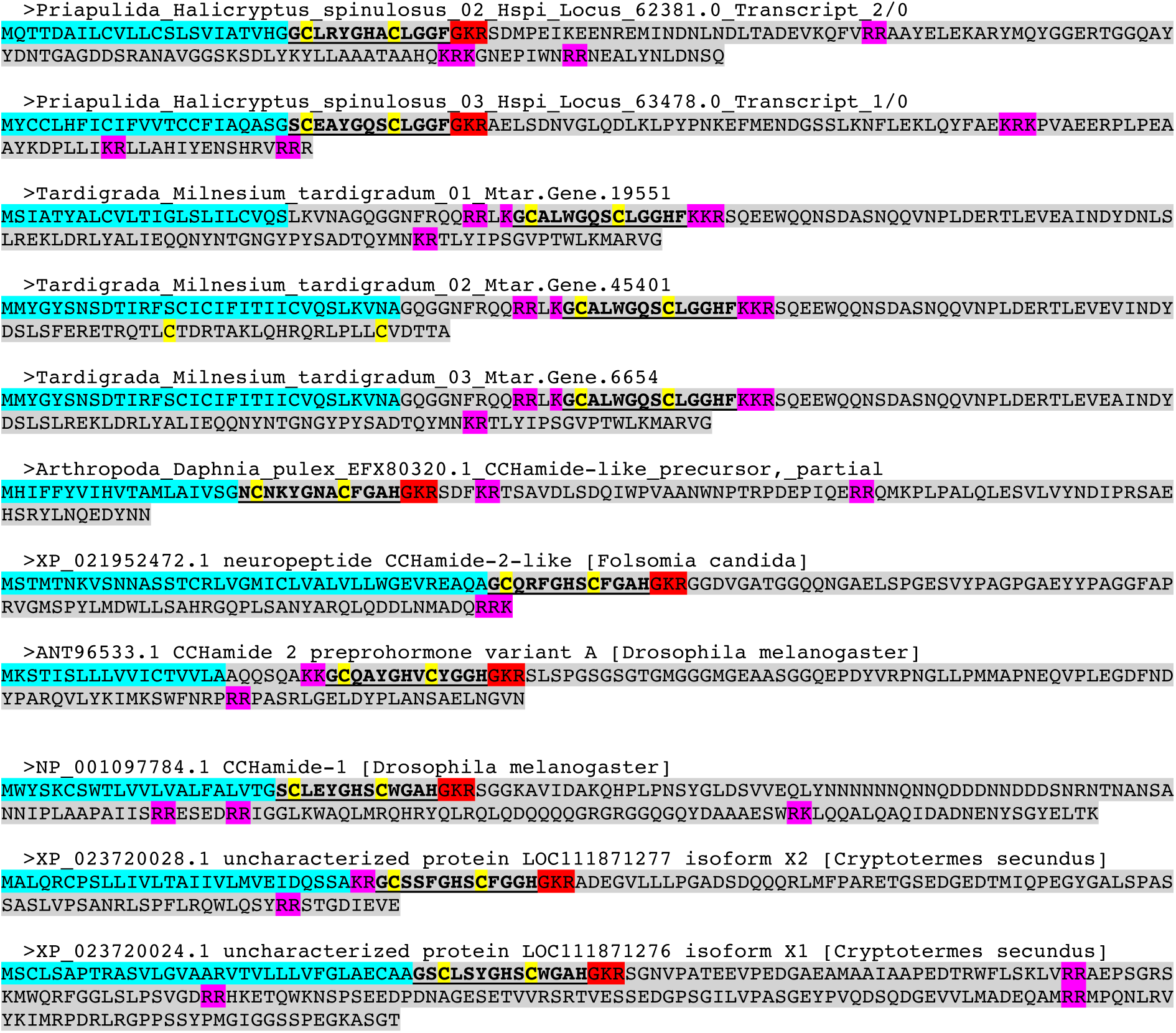

